# Multple-trait Bayesian Regression Methods with Mixture Priors for Genomic Prediction

**DOI:** 10.1101/102962

**Authors:** Hao Cheng, Kadir Kizilkaya, Jian Zeng, Dorian Garrick, Rohan Fernando

## Abstract

Bayesian multiple-regression methods incorporating different mixture priors for marker effects are widely used in genomic prediction. Improvement in prediction accuracies from using those methods, such as BayesB, BayesC and BayesC*π*, have been shown in single-trait analyses with both simulated data and real data. These methods have been extended to multi-trait analyses, but only under a specific limited circumstance that assumes a locus affects all the traits or none of them. In this paper, we develop and implement the most general multi-trait BayesCΠ and BayesB methods allowing a broader range of mixture priors. Further, we compare them to single-trait methods and the “restricted” multi-trait formulation using real data. In those data analyses, significant higher prediction accuracies were sometimes observed from these new broad-based multi-trait Bayesian multiple-regression methods. The software tool JWAS offers routines to perform the analyses.

## Introduction

Genomic prediction was proposed by Meuwissen et al. (Meuwissen et al. 2001) to incorporate whole-genome data into genetic evaluation. In genomic prediction, all the marker or haplotype effects are estimated simultaneously, and these estimates can then be used to predict breeding values of individuals not in the training population used to estimate the effects.

Bayesian multiple-regression methods incorporating mixture priors for marker effects are widely used in genomic prediction. For example, BayesB with locus specific variances accommodates models where markers have zero effects with probability *π* (Meuwissen et al. 2001; Cheng et al. 2015). Another mixture model, BayesC, assumes a common locus variance for all markers, and its extension known as BayesC*π* further treats *π* as an unknown parameter with a uniform prior distribution (Habier et al. 2011).

Bayesian multiple-regression methods were first proposed for single-trait analyses but have been extended to some particular forms of multi-trait analyses (Calus and Veerkamp 2011; Jia and Jannink 2012). Those extensions have pertained to a particular, somewhat restrictive mixture model. The “restricted” multi-trait BayesCΠ presented by Jia et al. (Jia and Jannink 2012) assumes a variant affects none of the traits or has simultaneous effects on all traits. This assumption of genetic architecture in that multi-trait BayesCΠ circumstance is violated if some loci have no effect on at least one of the traits while having an effect on at least one of the other traits.

In this paper, we present a more general class of multi-trait BayesCΠ and BayesB methods for which the previous multi-trait model is a special case. The new methods are compared to the previous multi-trait methods and to single-trait methods with real data.

## Materials and Methods

### Multi-trait Marker Effects Model

For simplicity and without loss of generality, we will assume a general mean as the only fixed effect, and write the multi-trait model for individual *i* from *n* genotyped individuals as 

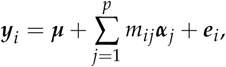

where ***y***_*i*_ is a vector of phenotypes of t traits for individual *i*, ***μ*** is a vector of overall means for *t* traits, *m_ij_* is the genotype covariateat locus *j* for individual *i, p* is the number of genotyped loci, ***α***_*j*_ is a vector of allele substitution effects of *t* traits for locus *j*, and ***e***_i_ is a vector of random residuals of *t* traits for individual *i*. The fixed effects, or general mean in this case, are assigned flat priors. The residuals, ***e***_*i*_, are a priori assumed to be independently and identically distributed multivariate normal vectors with null mean and covariance matrix ***R***, which in turn is assumed to have an inverse Wishart prior distribution, *W_t_*^-1^ (**S**_*e*_, *v_e_*).

## Multi-trait BayesCΠ model

### Priors for marker effects

The prior for *α_jk_*, the allele substitution or marker effect of trait *k* for locus *j*, is a mixture with a point mass at zero and a univariate normal distribution conditional on *sσ_k_*^2^: 

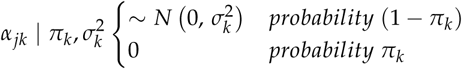

and the covariance between effects for traits *k* and *k’* at the same locus, i.e., *α_jk_* and *α_jk’_* is 

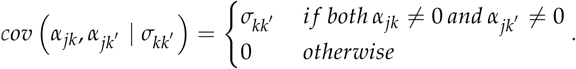

Employing the concept of data augmentation, the vector of marker effects at a particular locus ***α***_*j*_ can be written as ***α***_*j*_ = ***D***_*j*_***β***_*j*_, where ***D***_*j*_ is a diagonal matrix with elements *diag*(**D**_*j*_) = ***δ***_*j*_ = (*δ*_*j*1_, *δ*_*j*2_, *δ*_*j3*_ …), where *δ*_*jk*_ is an indicator variable indicating whether the marker effect of locus *j* for trait *k* is zero or non-zero, and ***β***_*j*_ follows a multivariate normal distribution with null mean and covariance matri 
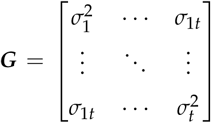
. The covariance matrix ***G*** is a priori assumed to follow an inverse Wishart distribution, *W_t_*^-1^(***S**_β_, v_β_*). Thus the multi-trait BayesCΠ model with data augmentation is written as 

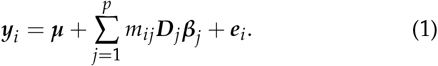

In the most general case, any marker effect might be zero for any possible combination of *t* traits resulting in 2^*t*^ possible combinations of ***δ***_*j*_. For example, in a *t*=2 trait model, there are 2^*t*^ = 4 combinations of ***δ***_*j*_, namely ***δ***_1_ = (0, 0), ***δ***_2_ = (0, 1), ***δ***_3_ = (1, 0), ***δ***_4_ = (1, 1). In the special case of this model described by (Jia and Jannink 2012), only ***δ***_1_ = (0, 0) and ***δ***_4_ = (1, 1) have non-zero probability. Suppose in general we use numerical labels “1”, “2”,…, “l” for the 2^*t*^ possible outcomes for ***δ***_*j*_, then the prior for ***δ***_*j*_ is a categorical distribution 

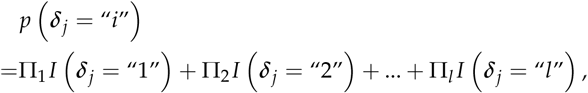

where Π_*i*_ is the probability that the vector *δ_j_* = “*i*” 
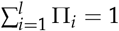
.

A Dirichlet distribution with all parameters equal to one, i.e., a uniform distribution, can be used for the prior for **Π** = (Π_1_, Π_2_, …, Π_*l*_). As shown below, a Gibbs sampler can be used to draw samples for all the parameters in this model.

### *Gibbs sampler I for multi-trait BayesC*Π

Suppose the prior for *δ*_*j*_ is a categorical distribution whose support is for all 2^*t*^ possible outcomes of ***δ***_*j*_. For convenience, from now on let “1” denote trait *k* and “2” the other *t* – 1 traits. In our sampling scheme, *β*_*j*1_ and *δ*_*j*1_ are sampled from their joint full conditional distributions, which can be written as the product of the full conditional distribution of *β*_*j*1_ given *δ*_*j*1_ and the marginal full conditional distribution of *δ*_*j*1_. Let ***θ*** denote all other parameters except *δ*_*j*1_ and *β*_*j*1_, then our sampling scheme can be written as 

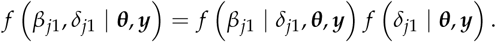

The full conditional distributions of *β*_*j*1_, *δ*_*j*1_, **Π, *G*** and ***R*** for Gibbs sampler **I**, which were derived in the Appendix, are given below.

The full conditional distributions of *β*_*j*1_ is 

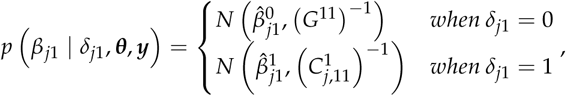

with 

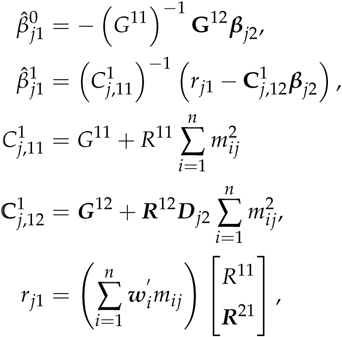

where 
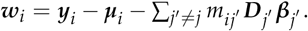

The marginal full conditional probability of *δ*_*j*1_ = 1 is 

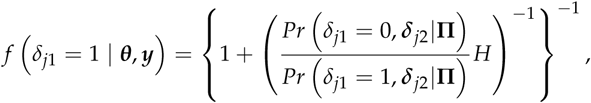

where 
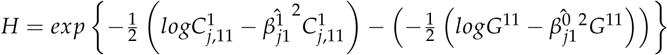
.

The full conditional distribution for **Π** can be written as 

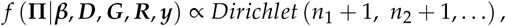

where *n_i_* is the number of markers with ***δ***_*j*_ = “*i*”.

The full conditional distributions for ***R***, the covariance matrix for residuals, is an inverse Wishart distribution, 
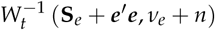
, where ***e*** is the *n* × *t* matrix for residuals with the *i*th row as ***e***_*i*_^′^. The full conditional distribution for ***G***, the covariance matrix for *β*_*j*_, is an inverse Wishart distribution, 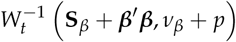
, where ***β*** is the *p* × *t* matrix with the *i*th row as ***β***_*i*_^′^.

**Figure 1.**
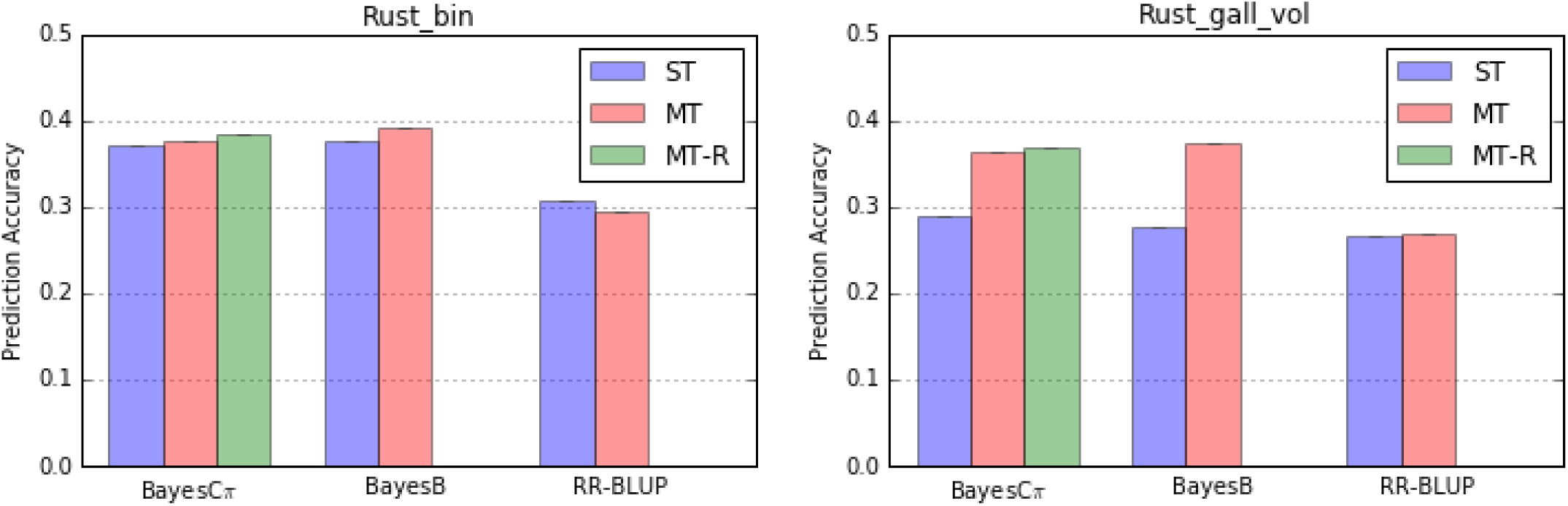
Comparison of single-trait and multi-trait methods for Rust_bin and Rust_gall_vol traits.

### *Gibbs sampler II for multi-trait BayesC*Π

The Gibbs sampler above requires that all 2^*t*^ outcomes for ***δ***_*j*_ have non-zero prior probabilities, i.e. none of Π_*i*_ can be zero. If some Π_*i*_ are zero, the markov chain generated from Gibbs sampler I may not be irreducible. Another more general Gibbs sampler that does not require all Π_*i*_ to be non-zero is proposed below.

The full conditional distributions of ***β***_*j*_, ***δ***_*j*_, **Π**, **G**, **R** for Gibbs sampler II, which were derived in the Appendix, are given below.

Let ***θ*** denote all other parameters except ***β***_*j*_ and ***δ***_*j*_, then our sampling scheme can be written as 

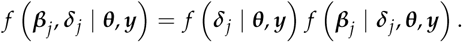

The full conditional distribution of ***β***_*j*_ is 

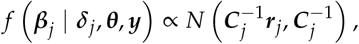

where
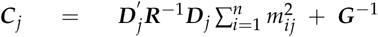
and 
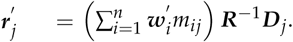

The marginal full conditional probability of ***δ***_*j*_ = “*i*” is 

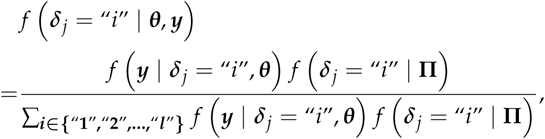

where 

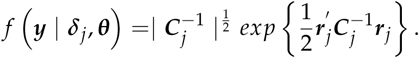

This Gibbs sampler can accommodate the restricted multi-trait BayesCΠ that was proposed by Jia et al. (Jia and Jannink 2012), which only allows ***δ***_*j*_ to be a vector of all ones or a vector of all zeros.

### Multi-trait BayesB Model

The multi-trait BayesCΠ model proposed above can be modified to accommodate the multi-trait BayesB model. Model equation (1) can also be used for the multi-trait BayesB method. The differences in multi-trait BayesB method is that the prior for ***β***_*j*_ is a multivariate normal distribution with null mean and locus-specific covariance matrix **G**_*j*_. The locus-specific covariance matrix ***G***_*j*_ is a priori assumed to follow an inverse Wishart distribution, 
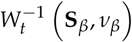

.

The derivations of the full conditional distributions of parameters of interest for Gibbs samplers are shown in the Appendix. In the multi-trait BayesB model, the full conditional distributions for all parameters except ***G***_*j*_ are similar to the multi-trait BayesCΠ model. The full conditional distribution for ***G***_*j*_, the covariance matrix for *β*_*j*_, is a inverse Wishart distribution, 
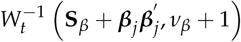
.

## Data analyses

Published genotypic and deregressed phenotypic data for Loblolly Pine (Pinus Taeda L.) were used (Resende et al. 2012). Two disease traits, namely Rust_bin and Rust_gall_vol were analyzed. The reported heritability was 0.21 for Rust_bin and 0.12 for Rust_gall_vol. Loci with missing genotypes were imputed as the mean of the observed genotype covariates at that locus and loci with a missing rate >50% were excluded. After these quality control edits, 4,828 SNPs on 807 individuals with phenotypes and genotypes on both traits remained.

Prediction accuracy was calculated as the correlation between the vector of deregressed phenotypes and the vector of estimated breeding values. Cross-validation using 10-folds formed the basis for comparing our general multi-trait BayesCΠ model (MT-BayesCΠ) to a similar model where the prior for ***β**_**j**_* is a multivariate normal rather than a mixture of multivariate normals (MT-BayesC0), the restricted multi-trait BayesCΠ proposed by Jia at al. (MT-BayesCΠ-R), multi-trait BayesB with known Π (MT-BayesB) and the usual single trait formulations of the mixture models (ST-BayesC0, ST-BayesC*π*, ST-BayesB). The constant Π used in BayesB were estimated using BayesCΠ methods. All analyses were performed using JWAS (Cheng et al. 2016), a publicly-available package for single-trait and multi-trait whole-genome analyses written in the freely-available Julia language. Since BayesC0 is equivalent to random regression best linear unbiased prediction (RR-BLUP), ST-BayesC0 and MT-BayesC0 are denoted as ST-RR-BLUP and MT-RR-BLUP below. The prior for the residual covariance matrix **R** in all multi-trait methods was an inverse Wishart distribution, 
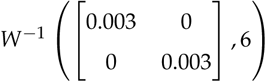
 or which the mean of R is
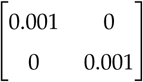
. The prior for the marker effects covariance matrix **G** in MT-BayesCΠ and MT-BayesCΠ-R was an inverse Wishart distribution, 
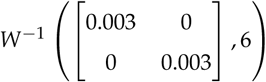
, for which the mean of ***G*** was 
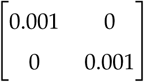
. The priors for the residual variance and marker effects variance in single-trait analyses were a scaled inverted chi-squared distribution with scale parameter S^2^ = 0.0005 and degrees of freedom *v* = 4, for which the mean of the prior was also 0.001. Marker effect variances estimated from BayesCΠ were used to construct the priors for marker effect variances in the BayesB methods.

## Results

The prediction accuracies from all methods for Rust_bin and Rust_gall_vol are in figure 1. The prediction accuracies from all single-trait analyses using JWAS are similar to those in (Resende et al. 2012). ST-BayesC*π* showed higher prediction accuracies than ST-RR-BLUP for both traits (Resende et al. 2012). The prediction accuracies from ST-BayesB were similar to those from ST-BayesC*π*, when both marker effect variances and *π* estimated from ST-BayesC*π* were used in ST-BayesB.

The analyses of Rust_bin exhibited no significant difference between multi-trait and single-trait analyses within each method (ST-RR-BLUP versus MT-RR-BLUP; ST-BayesC*π* versus MT-BayesCΠ; ST-BayesC*π* versus MT-BayesCΠ-R; ST-BayesB versus MT-BayesB).

In contrast, analyses for the lower heritability Rust_gall_vol with MT-BayesCΠ showed significantly higher accuracies than ST-BayesC*π*. MT-BayesCΠ and MT-BayesCΠ-R showed similar prediction accuracies. The posterior means of P for both methods were shown in table 1. The performance of MT-BayesB were similar to MT-BayesCΠ, when both marker effect variances and P estimated from MT-BayesCΠ were used. Similar prediction accuracies were observed in MT-RR-BLUP and ST-RR-BLUP for trait Rust_gall_vol.

## Discussion

In the single trait analyses, accuracies from ST-BayesC*π* and ST-BayesB were higher than those from ST-RR-BLUP, suggesting that these two traits are influenced by a few QTL with large effects. The effect of genetic architecture on the performance of multi-trait analyses has been studied in previous simulation analyses (Jia and Jannink 2012). Using simulated data they found that multi-trait Bayesian variable selection methods outperform multi-trait RR-BLUP in the presence of major QTL. This observation was confirmed in our real data analyses that MT-BayesCΠ and MT-BayesB outperformed MT-RR-BLUP for both traits.

Significant differences between multi-trait and single-trait analyses were only observed for Rust_gall_vol within BayesC*π* and BayesB methods (MT-BayesCΠ versus ST-BayesC*π*; MT-BayesB versus ST-BayesB). MT-BayesCΠ and MT-BayesCΠ-R outperformed ST-BayesC*π* for Rust_gall_vol, and the accuracy gain was 26% (from 0.287 to 0.364). The lower-heritability trait Rust_gall_vol may borrow information from the other correlated trait Rust_bin. Thus higher prediction accuracy from MT-BayesCΠ were observed in trait Rust_gall_vol instead of Rust_bin. Results in (Jia and Jannink 2012) showed no difference between MT-BayesCΠ-R and ST-BayesC*π* because a reduced marker panel (500 markers) was used. The performance of MT-BayesB was similar to MT-BayesCΠ, when both marker effect variances and P estimated from MT-BayesCΠ were used. Further analyses may be required to study the effects of priors in MT-BayesB.

The fact that RR-BLUP showed no improvement in multi-trait analyses suggested that benefits from MT-BayesCΠ may caused by the estimation of hyper-parameter Π. In the MT-BayesCΠ, the mean of the posterior probability that a marker has a null effect on Rust_gall_vol was about 0.97, calculated as the summation of posterior mean of P for categories (0, 0) and (1, 0). The posterior mean of *π*, the probability that a marker has a null effect, in ST-BayesC*π* for Rust_gall_vol was 0.74, different from the equivalent value, 0.97, in MT-BayesCΠ showed above. Thus ST-BayesC*π* with constant *π*, equal to 0.97, were performed. Prediction accuracies from ST-BayesC*π* with constant *π* = 0.97 was 0.361, which was similar to the accuracies from MT-BayesCΠ. This suggests that high-heritability traits may help with variable selection in correlated low-heritability traits.

The difference between MT-BayesCΠ and MT-BayesCΠ-R is that MT-BayesCΠ-R assumes a locus has an effect on all traits or none of them. This assumption of genetic architecture is always violated. MT-BayesCΠ and MT-BayesCΠ-R, however, showed similar prediction accuracies. This can be explained by the estimation of Π in MT-BayesCΠ and MT-BayesCΠ-R in table 1. The posterior probability means for (0, 1) and (1, 0) were almost zero in MT-BayesCΠ and for (0, 0) and (1, 1) are similar in MT-BayesCΠ and MT-BayesCΠ-R, suggesting that the assumption of genetic architecture for MT-BayesCΠ-R is valid for these two traits.

In practice, genetic variances from previous conventional analyses are always used to construct priors for marker effect variances. For single trait analyses, under some assumptions, it can be shown that the marker effect variance 
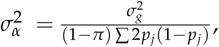
 where *σ_g_*^2^ is the genetic variance, *p_j_* is the allele frequency for locus *j* and *π* is the probability that a marker has a null effect. Following similar strategies, the marker effect covariance matrix **G** in two-trait analyses can be obtained as 

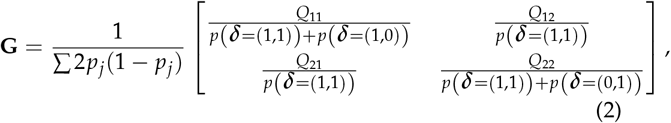

where 
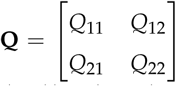
 the genetic covariance matrix and 
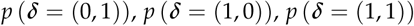
 are the probability a marker has null effects on the first trait but not the second trait, on the second trait but not the first trait and on no traits. Thus the probability that a marker has an effect on the first trait can be obtained as 
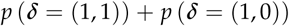
, which is the denominator of the upper left element in (2). This strategy relating genetic covariance matrix to marker effect covariance matrix can also be used for analyses with more than two traits. Note that positive definite matrix **Q** may result in negative definite matrix **G** using (2), especially when the prior for the probability a marker has null effects violates the truth.

**Table 1.**
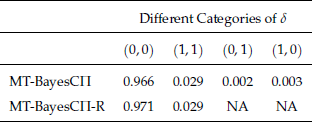
Estimation of π for alternative multi-trait BayesCΠ methods. Posterior mean of Π were given for different categories of *δ*. Different categories of *δ* are denoted as(*k*_1_, *k*_2_), where *k*_1_ = 0 if a marker has a null effect on Rust_bin, otherwise *k*_1_ = 1, and similarly for *k*_2_ representing sampled effects for Rust_gall_vol. Combinations listed as NA do not exist in the restricted model.

## Appendix

### Gibbs sampler algorithm for multi-trait BayesCΠ

#### *Single-site Gibbs sampler for multi-trait BayesC*Π

The full conditional distribution of *β*_*j*1_ can be written as 

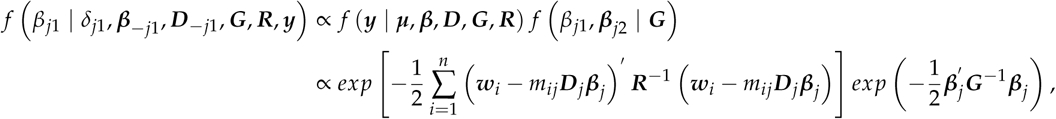

where 
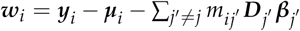
. Further, by dropping factors that do not involve *β*_*j*1_, 

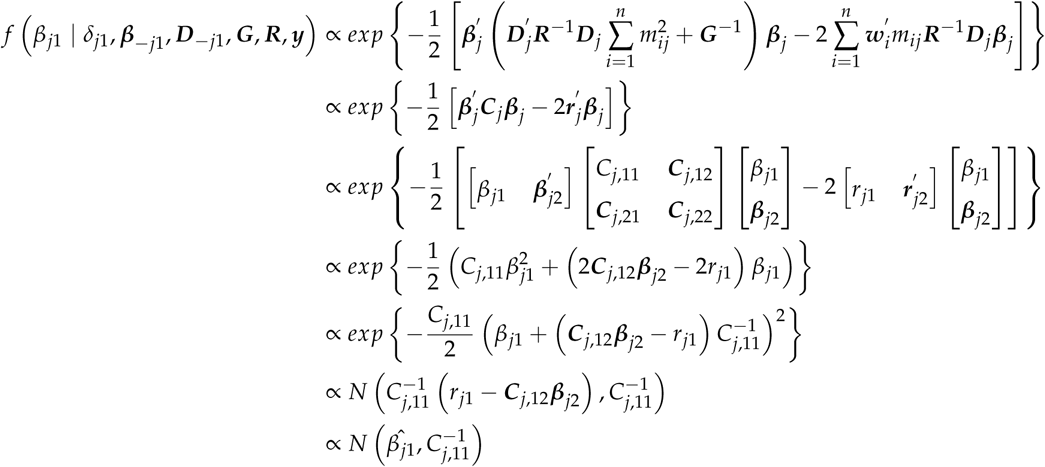

where 
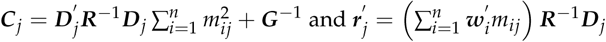
.

Note that when *δ*_*j*1_ = 0, 

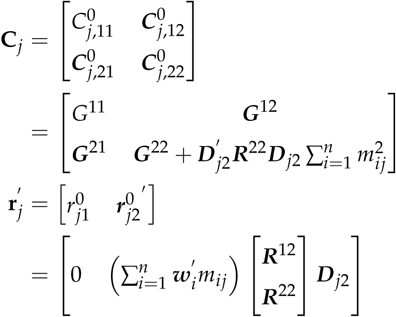

When *δ*_*j*1_ = 1, 

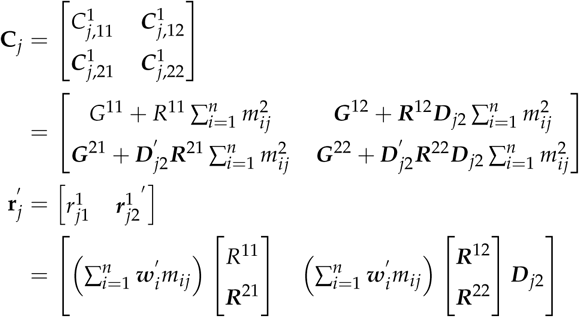

Thus when *δ*_*j*1_ = 0, the full conditional distribution of *β*_*j*1_ is 

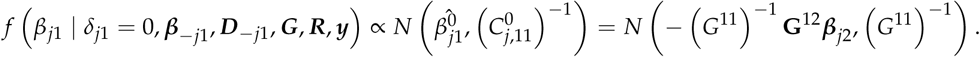

When *δ*_*j*1_ = 1, the full conditional distribution of *β*_*j*1_ becomes 

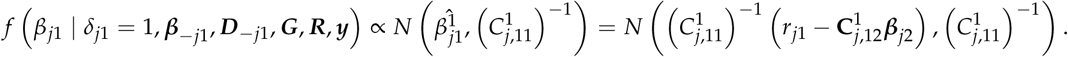

The marginal full conditional distribution of *δ*_*j*1_ can be written as 

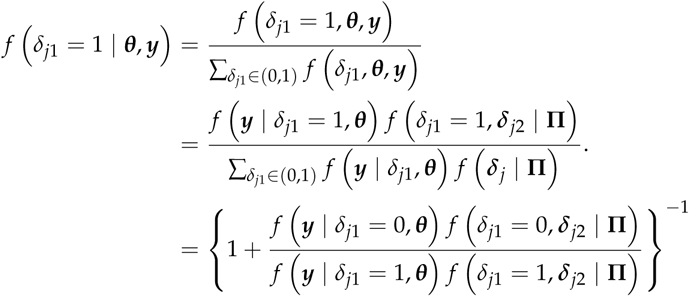

The factor 
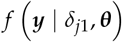
 can be written as 

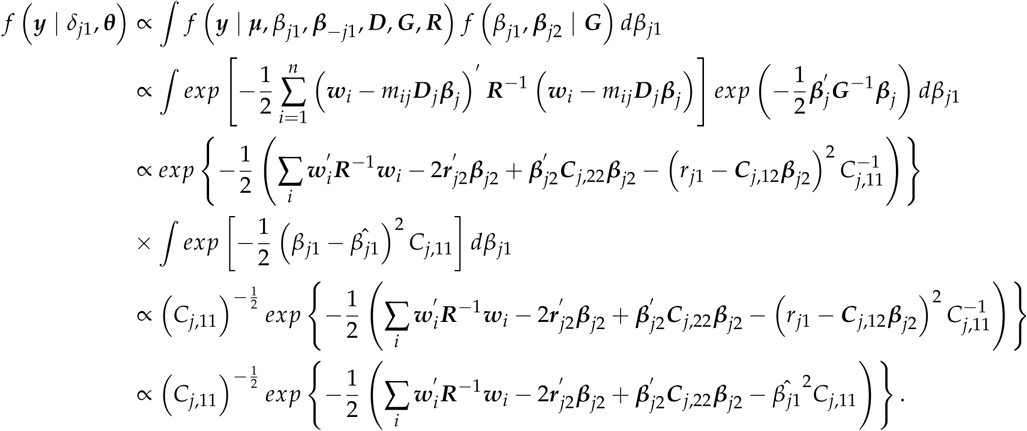

Note that 
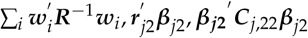
 are same when *δ*_*j*1_ = 0 or 1. Thus the ratio 
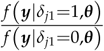
 becomes 

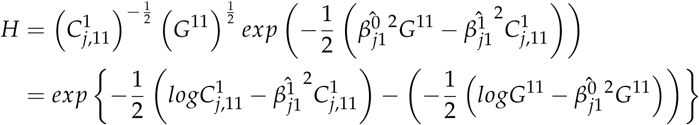

Thus the conditional probability of *δ*_*j*1_ = 1 is 

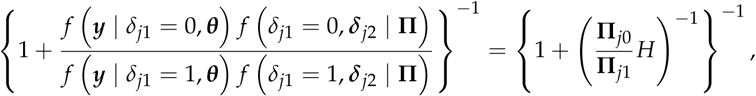

where 
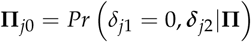
 and 
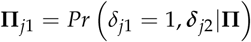

The full conditional distribution for **Π** can be written as 

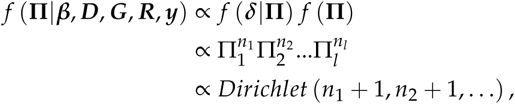

where *n_i_* is the number of markers with ***δ***_*j*_ = “*i*”.

#### *Joint Gibbs sampler for multi-trait BayesC*Π

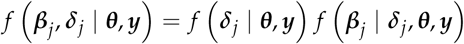

The marginal full conditional distribution of ***δ***_*j*_ can be written as 

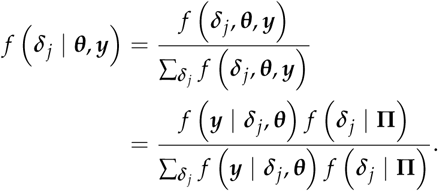

Denote 
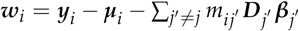
, then 

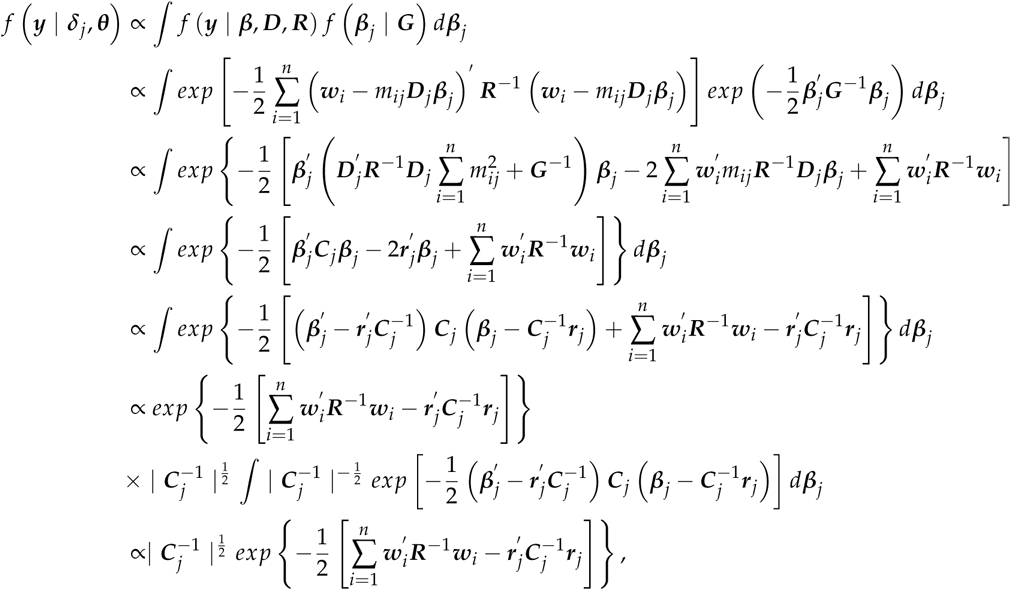

where 
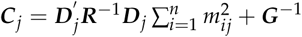
 and 
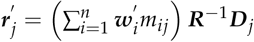
.

Note that 
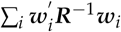
 is same for different ***δ***_*j*_. Thus the marginal full conditional distribution of ***δ***_*j*_ can be written as 

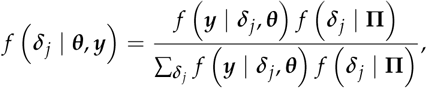

where 

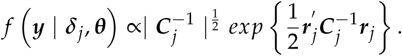

The full conditional distribution of ***β***_*j*_ is 

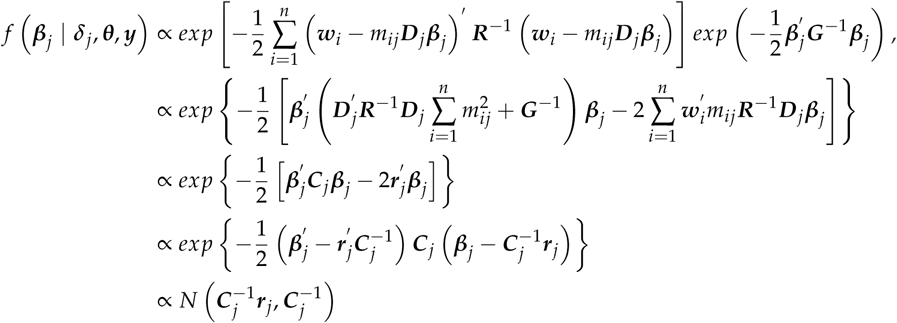

#### Gibbs sampler algorithm for multi-trait BayesB

##### Single-site Gibbs sampler for multi-trait BayesB

For convenience, from now on let “1” denote trait *k* and “2” the other traits. Thus, ***β***_*j*_ can be denoted as 
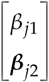
 and ***D***_*j*_ can be denoted as 
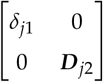
. The Gibbs sampler for *β*_*jk*_ and *δ*_*jk*_ is derived as below. In our sampling scheme, *β*_*j*1_ and *δ*_*j*1_ are sampled from their joint full conditional distributions, which can be written as the product of the full conditional distribution of *β*_*j*1_ given *δ*_*j*1_ and the marginal full conditional distribution of *δ*_*j*_. Let ***θ*** denote all other parameters except *δ*_*j*1_ and *β*_*j*1_, then our sampling scheme can be written as 

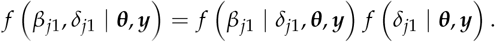

The full conditional distribution of *β*_*j*_ can be written as 

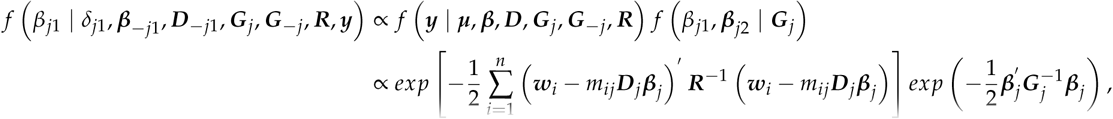

where 
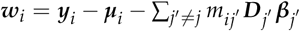
. Further, by dropping factors that do not involve *β*_*j*1_, 

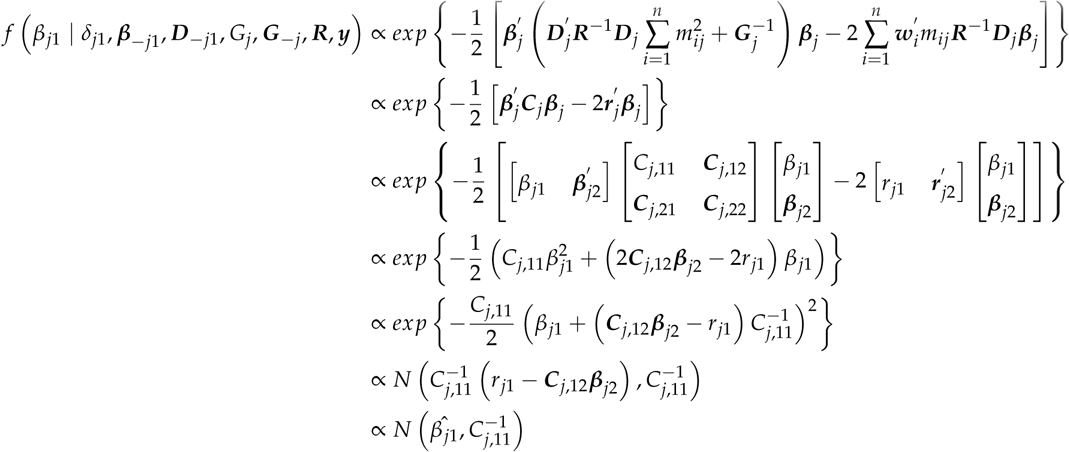

where 
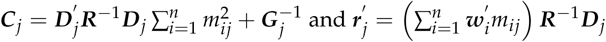

Note that when *δ*_*j*1_ = 0, 

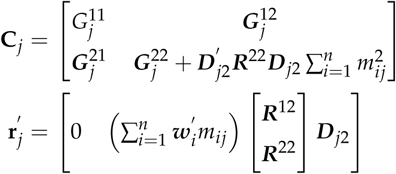

When *δ*_*j*1_ = 1, 

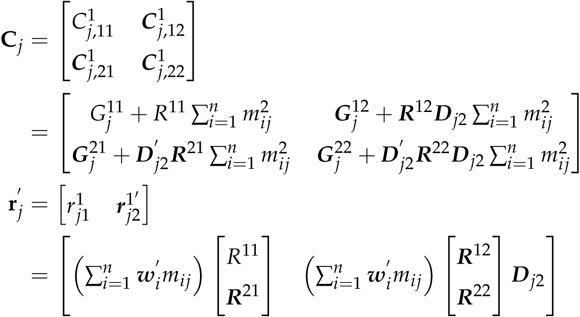

Thus when *δ*_*j*1_ = 0, the full conditional distribution of *β*_*j*1_ is 

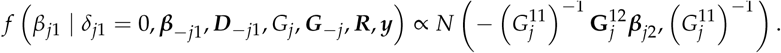

When *δ*_*j*1_ = 1, the full conditional distribution of *β*_*j*1_ becomes 

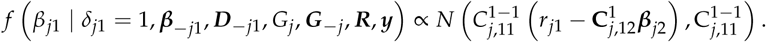

The marginal full conditional distribution of *δ*_*j*1_ can be written as 

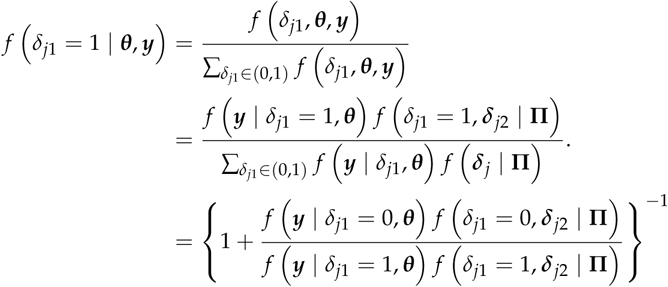

The factor 
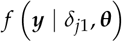
 can be written as 

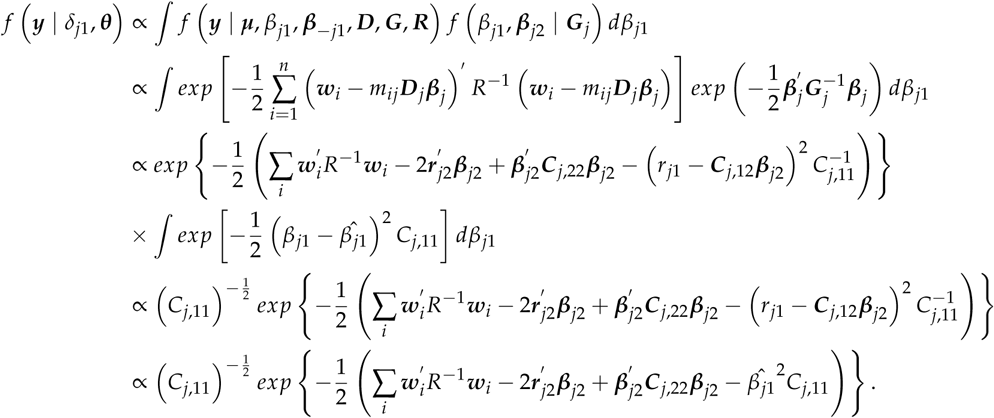

Note that 
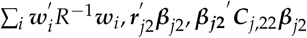
are same when *δ*_*j*1_ = 0 or 1. Thus the ratio 
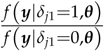
 becomes 

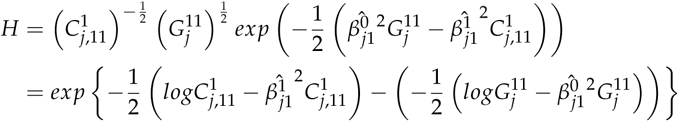

Thus the conditional probability of *δ*_*j*1_ = 1 is 

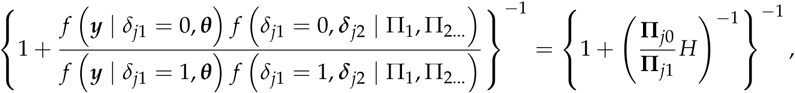

where 
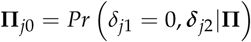
 and 
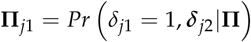
.

#### Joint Gibbs sampler for multi-trait BayesB

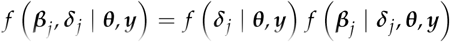

The marginal full conditional distribution of ***δ***_*j*_ can be written as 

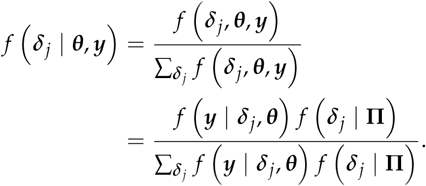

Denote 
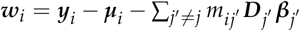
, then 

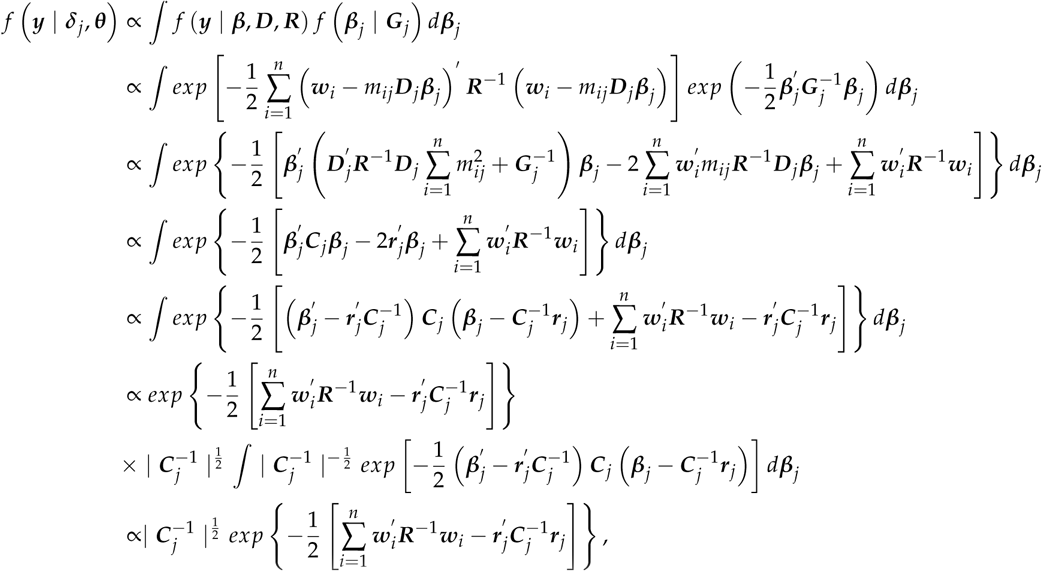

where 
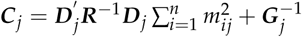
 and 
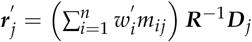
.

Note that 
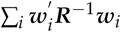
 is same for different ***δ***_*j*_. Thus the marginal full conditional distribution of ***δ***_*j*_ can be written as 

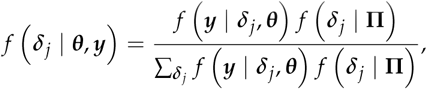

where 

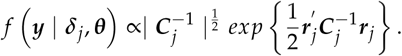

The full conditional distribution of ***β***_*j*_ is 

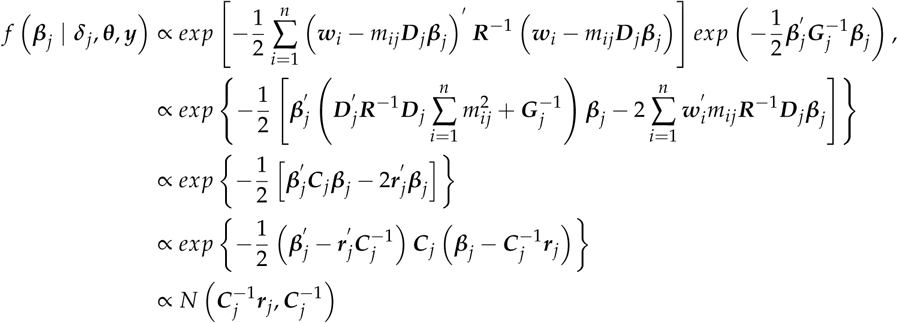

